# A split crRNA with CRISPR-Cas12a enables highly sensitive, selective, and multiplexed detection of RNA and DNA

**DOI:** 10.1101/2024.01.04.574169

**Authors:** Jie Qiao, Yichuan Chen, Xinping Wang, Baoxia He, Wenhao Yin, Yi Liu

## Abstract

The complete 40-nt CRISPR RNA (crRNA) of Cas12a can be artificially divided into two parts, including the 20-nt scaffold RNA with fixed sequences and the 20-nt spacer RNA with variable sequences, respectively. Herein, we found that Cas12a protein, scaffold RNA, and spacer RNA can be reassembled into an active ribonucleoprotein (RNP) which has a *trans*-cleavage activity comparable to that of wild-type Cas12a RNP. By leveraging such split CRISPR-Cas12a system (SCas12a), we devised fast fluorescence and lateral flow assays for highly sensitive, selective, and multiplexed detection of miRNAs without the need for reverse transcription and pre-amplification, achieving a limit of detection (LoD) of 10 fM. Additionally, our SCas12a assay enables detecting long-stranded RNA without secondary structure, as well as distinguishing mature miRNA from its precursor (pre-miRNA) that comprises the same sequence of miRNA. Beyond RNA detection, SCas12a outperforms wild-type Cas12a in specificity towards DNA point mutations. In combination with recombinase polymerase amplification (RPA), we set up a one-pot assay to detect attomolar concentrations of human papillomavirus (HPV) in patient samples. In conclusion, this work provides a simple, cost-effective, yet powerful SCas12a-based rapid nucleic acid detection platform in various diagnostic settings.

**Significance Statement:** Conventional Cas12a-based diagnostic methods cannot directly detect RNA targets. Here, we develop rapid fluorescence and lateral flow assays based on a split Cas12a system (called SCas12a), achieving amplification-free detection of RNA with high sensitivity and specificity. By supplying pooled activators, our method enables multiplexed detection of miRNA and DNA targets. In addition, SCas12a can discriminate mature miRNA from its pre-miRNA, which was achieved for the first time with CRISPR detection technology. Moreover, SCas12a outperforms wild-type Cas12a in specificity towards point mutation and can be combined with RPA to detect attomolar concentration of HPV in patient samples. Overall, this work offers a generic “split crRNA-activator” strategy for developing new CRISPR diagnostic tools.

## Introduction

Fast, accurate, and cost-effective nucleic acid testing (NAT) is vital for wide areas, including public health, disease diagnostics, environmental sciences, food safety, etc. Over the past decades, quantitative polymerase chain reaction (qPCR) has been considered as a gold standard technique for NAT(1, 2). However, the qPCR-based diagnostic methods require expensive instruments and time-consuming procedures, significantly limiting their applications in the field of point-of-care testing (POCT). To overcome these shortcomings, the CRISPR (clustered regularly interspaced short palindromic repeats)-Cas enzyme (Cas12a (3, 4), Cas12b (5), Cas13a (6), Cas14 (7) etc.) based analytical methods (8, 9) have been rapidly developed in recent years. Among the discovered CRISPR-Cas enzymes, CRISPR-Cas12a exhibits a unique ability that can indiscriminately cut any nearby non-specific single strand DNA (ssDNA) molecules after target-specific recognition and cleavage of DNA (10). By leveraging such *trans*-cleavage catalytic property, a variety of diagnostic tools (e.g. DETECTR (3, 11) and HOLMES(4)) have been invented for ultrasensitive detection of DNA and RNA. Unlike directly detecting DNA, the RNA substrates must be firstly converted to DNA by reverse transcription (12, 13) before they can be recognized by Cas12a. If the detected object is miRNA (14), a short RNA of ∼ 20-22 nucleotides, it is more difficult to obtain high-quality DNAs by reverse transcription. Additional sample amplification steps are often required, resulting in long experiment times (∼ 2 hours) and significantly increasing the risk of aerosol contamination. Although other CRISPR enzymes such as CRISPR-Cas13a enable direct detection of miRNA, they remained challenging to eliminate the interferences of longer precursor RNAs (pre-miRNAs). The limit of detection (LoD) of these Cas13a-based methods (e.g. SHERLOCK (15, 16)) is usually higher than 10 pM towards miRNA (17), which is not good enough for several clinical applications. Compared to Cas12a that uses ssDNA probes, Cas13a uses RNA probes (18) that are more expensive and easily degradable, leading to a higher risk of false positive results and higher testing costs. Taken together, it is urgent to develop new Cas12a-based diagnostic tools to address these issues.

Recently, Jain et al. developed a diagnostic method that, for the first time, enables direct detection of RNA by Cas12a (19). By supplying a short ssDNA or a PAM-containing dsDNA at the seed region of the crRNA, the method detected RNA substrates at the 3’-end of the crRNA. However, this “split-activator combination” strategy just partially regained the diminished activity of Cas12a ribonucleoprotein (RNP), obtaining a high LoD of 100-700 pM against different RNA targets. Therefore, Cas12a assay still needs to be significantly improved for RNA detection. In our recent work (20), we happened to find that the Cas12a protein can bind tightly to a truncated fragment of crRNA (called scaffold RNA) with a 20-nt fixed sequence: 5’-AAUUUCUACUAAGUGUAGAU-3’. This “Cas12a-scaffold RNA” complex has also recently been reported by other groups (21, 22). Compared to wild-type Cas12a RNP (Figure 1a) that has a 40-nt intact crRNA, this impaired Cas12a RNP did not exhibit any nuclease activity, possibly due to a lack of spacer RNA. Inspired by these findings, here we designed a new split Cas12a system (SCas12a, Figure 1b), consisting of Cas12a protein, scaffold RNA, and spacer RNA.

**Figure 1.**
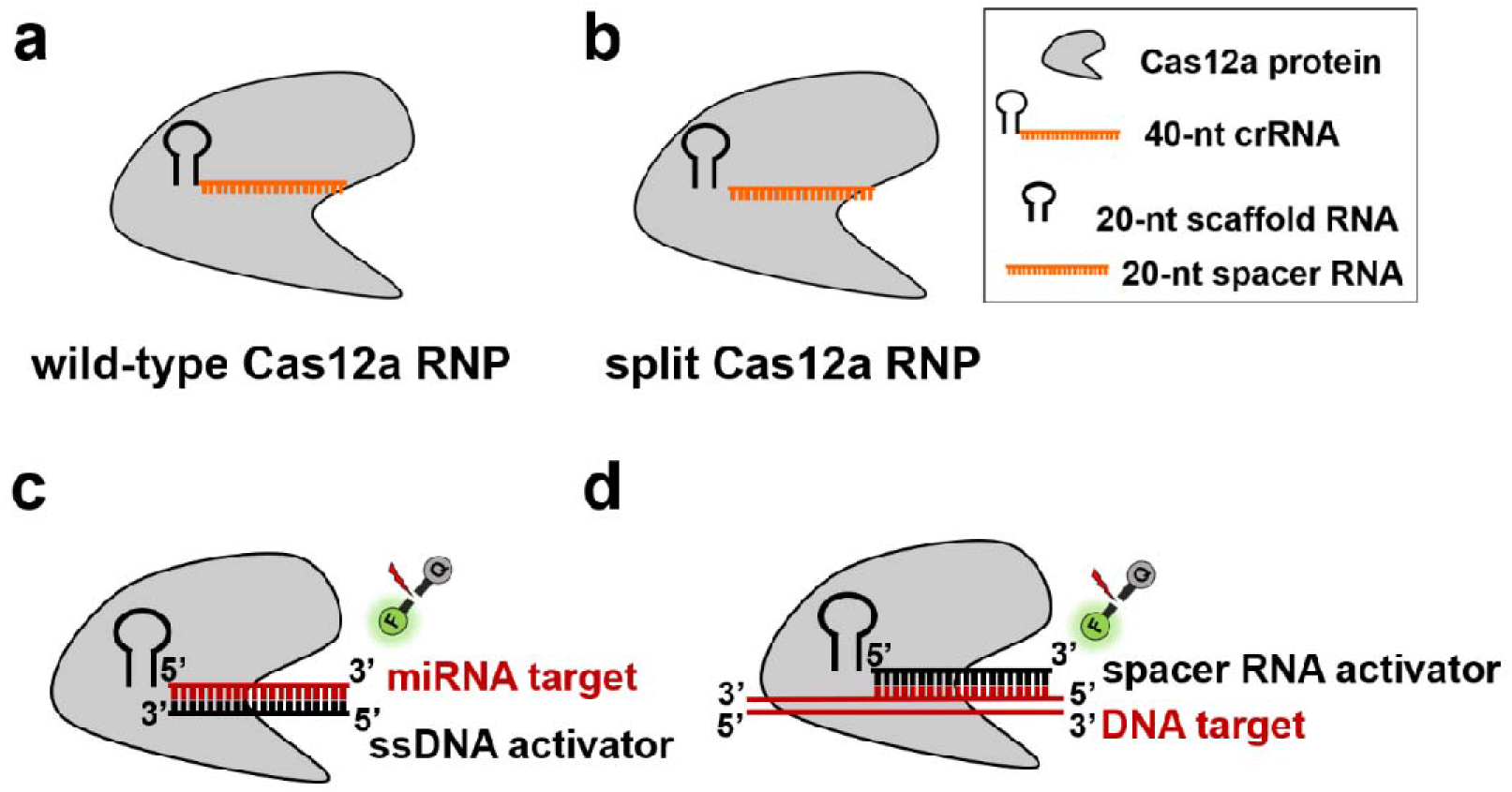
Schematic of the split Cas12a-based assay for direct detection of miRNA and DNA. (a) Schematic diagram of the wild-type Cas12a RNP. (b) Schematic diagram of the split Cas12a RNP. (c) The working mechanism of the amplification-free fluorescence assay developed for highly sensitive and selective detection of miRNA. (d) The working mechanism of the fluorescence assay for highly sensitive and selective detection of DNA in this study.

Here, we showed that the SCas12a RNP has a *trans*-cleavage activity comparable to that of wild-type Cas12a RNP. By leveraging SCas12a, we first developed a fluorescence assay for highly sensitive, selective, and multiplexed detection of miRNAs without additional reverse transcription and pre-amplification, achieving a limit of detection (LoD) of 100 fM. Next, we devised a lateral flow assay for rapid miRNA detection with a lower LoD of 10 fM. In addition, our method showed the ability to detect long-length RNA without secondary structure. Most importantly, our Cas12a assay was able to discriminate between mature miRNA and pre-miRNA that comprise the same sequence of miRNA, while Cas13a-based diagnostics cannot do that. Finally, we showed that SCas12a outperforms wild-type Cas12a in specificity towards DNA point-mutations and enables one-pot detection of attomolar concentrations of human papillomavirus (HPV) in combination with recombinase polymerase amplification (RPA). Together, we established a SCas12a-based diagnostic method for simple, rapid, cost-effective, and sensitive detection of RNA and DNA.

## Results

### Proof-of-principle studies for the development of SCas12a-based diagnostic method

Figure 1 illustrates the working mechanism of our split Cas12a system. To obtain the designed SCas12a RNP (Figure 1b), we firstly mixed the purified recombinant *Acidaminococcus* sp. Cas12a (AsCas12a) protein (Figure S1) with synthesized scaffold RNA and spacer RNA at a ratio of 1: 2: 2 *in vitro*. Next, we measured the on-target cleavage (*cis*-cleavage, Figure 2a) activity of this variant in the presence of targeted dsDNA plasmids. We also examined the collateral cleavage (*trans*-cleavage, Figure 2b-2c) ability of the variant by addition of the complementary dsDNA or ssDNA substrates, as well as non-specific ssDNA probes for cutting. The PAGE data revealed that nearly 100% *cis*- and *trans*-cleavage activities of the wild-type Cas12a RNP were recovered, which is consistent with a previous study (22). Encouraged by these critical results, we then performed proof-of-principle studies for the development of SCas12a-based diagnostic method.

**Figure 2.**
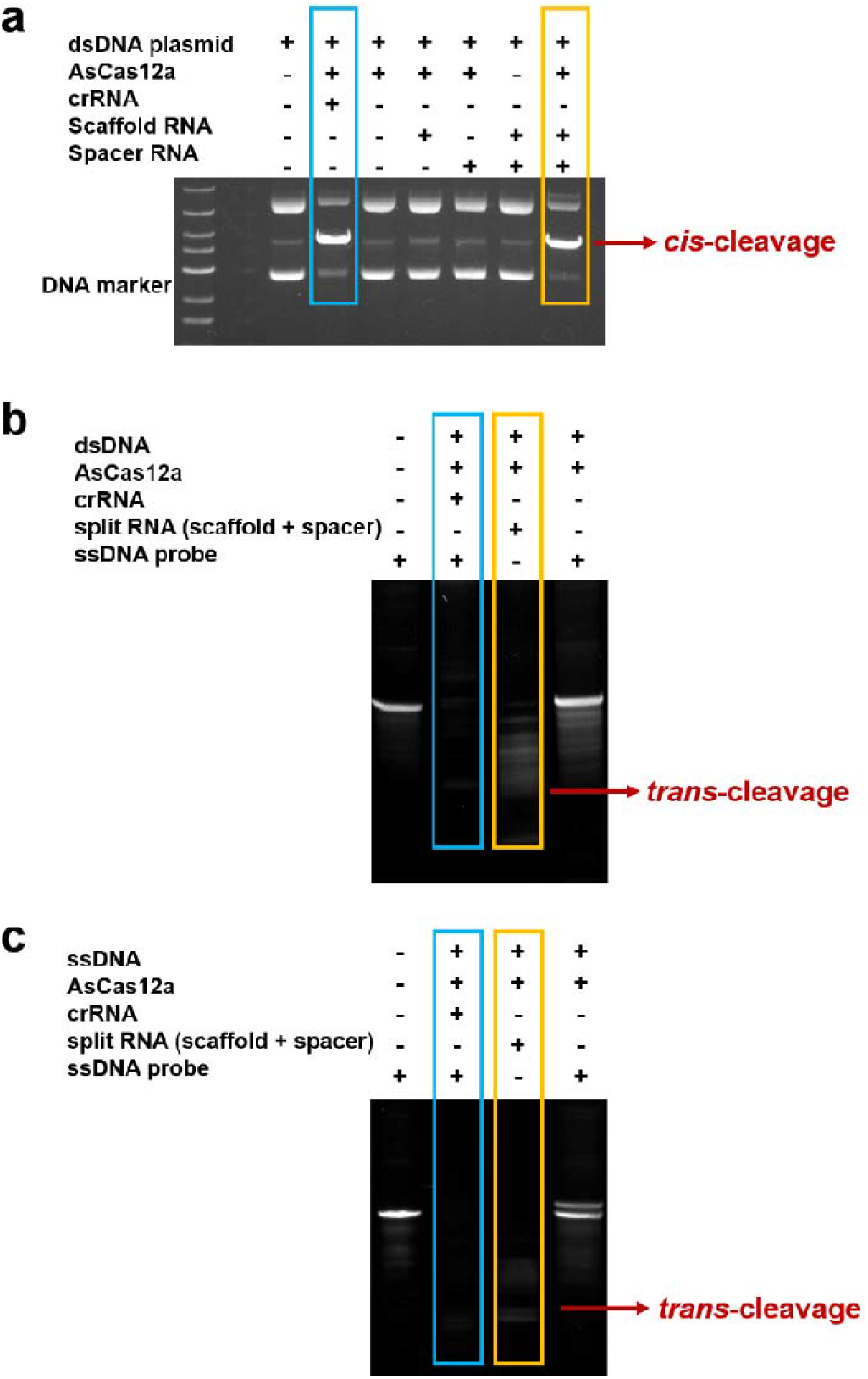
Comparison of the nuclease activities between the split Cas12a RNP and wild-type Cas12a RNP. (a) *Cis*-cleavage of target dsDNA plasmids by wild-type Cas12a RNP (blue) and split Cas12a RNP (orange). (b) *Trans*-cleavage of ssDNA probes by wild-type Cas12a RNP (blue) and split Cas12a RNP (orange) in the presence of complementary dsDNA substrates. (c) *Trans*-cleavage of ssDNA probes by wild-type Cas12a RNP (blue) and split Cas12a RNP (orange) in the presence of complementary ssDNA substrates. Reactions were incubated for 30 min at 37°C. The reactions contained 250 nM Cas12a, 500 nM crRNA, 500 nM scaffold RNA and 500 nM spacer RNA. For *cis*-cleavage, it contained (a) 330 ng dsDNA plasmids. For *trans*-cleavage, the reactions contained (a) 500 nM dsDNA or (b) 500 nM ssDNA activators, as well as 1000 nM ssDNA probes.

As shown in Figure 3a, a very strong “turn-on” fluorescence was generated only when Cas12a enzyme, scaffold RNA, spacer RNA, and targeting dsDNA all existed. The fluorescence intensity of SCas12a RNP was identical to that of wild-type Cas12a RNP. On the contrary, neither Cas12a nor Cas12a plus scaffold RNA unlocked collateral cleavage activity and almost no fluorescence signals were observed. Cas12a plus spacer RNA only partially activated *trans*-cleavage and a significantly reduced fluorescence signal was generated, which demonstrated that Cas12a protein, scaffold RNA and spacer RNA are all necessary to rebuilt the Cas12a RNP with full *trans*-cleavage activity. Moreover, the non-specific DNA substrates cannot initiate the *trans*-cleavage activity of SCas12a system, indicating the high specificity of our assay. Under the same reaction conditions, we further found that the 20-nt ssDNA target also activated *trans*-cleavage (Figure 3b). Finally, we carried out kinetic assays by adding spacer RNAs with different sequences (Table S1). All the reactions were completed within 10 minutes without sequence basis (Figure S2), indicating that our assay has the potential to become a universal NAT platform.

**Figure 3.**
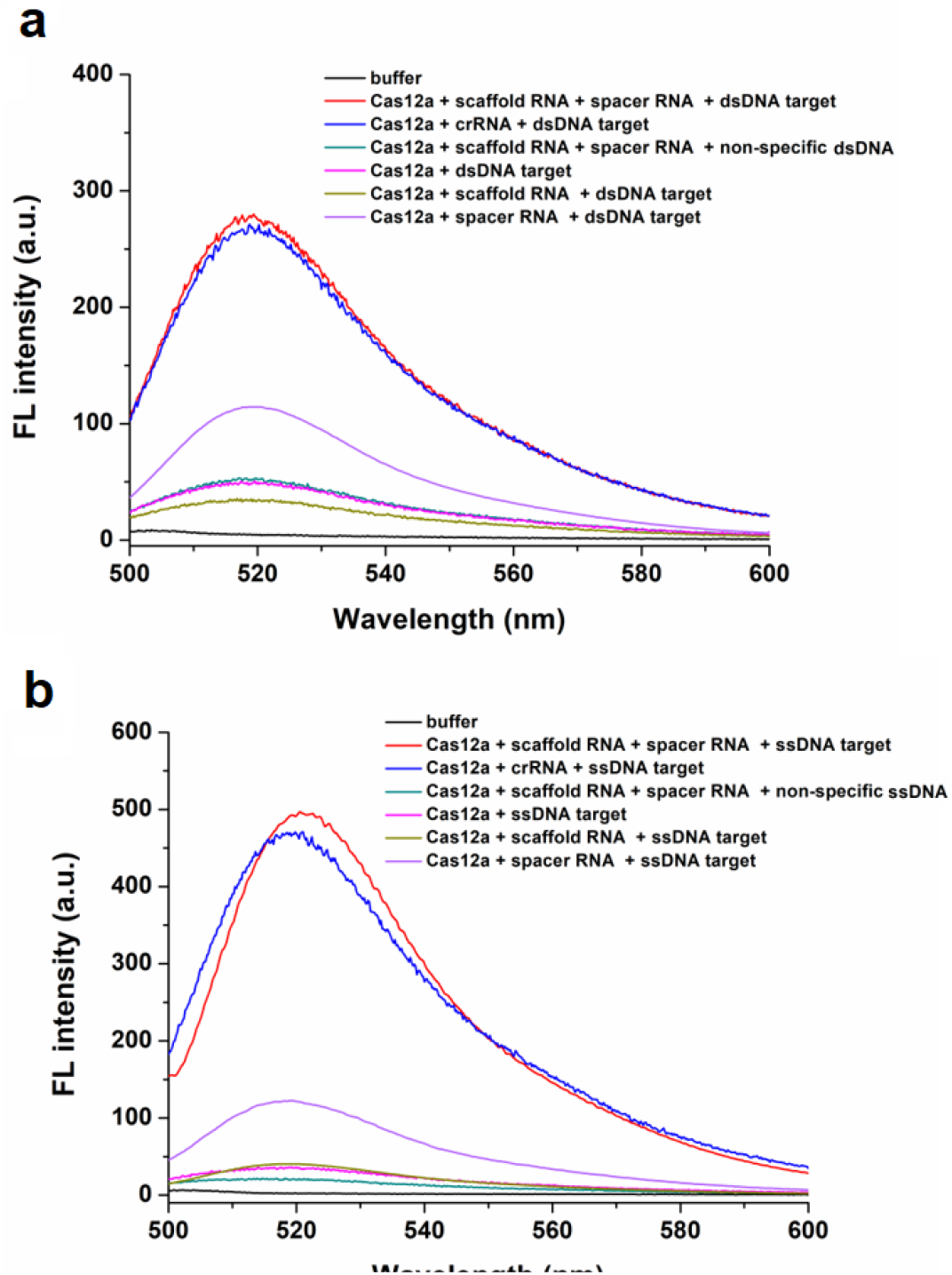
Proof-of-principle of the SCas12a-based fluorescence assay. (a) Detection of dsDNA by SCas12a or WT Cas12a. (b) Detection of ssDNA targets by SCa12a or WT Cas12a. Reactions were incubated for 30 min at 37°C. The reactions contained 250 nM Cas12a, 500 nM dsDNA target (a) or 500 nM ssDNA target (b), 500 nM crRNA, 500 nM scaffold RNA, 500 nM spacer RNA and 1000 nM fluorescence probe.

### Amplification-free miRNA detection by SCas12a-based assay

At first, we noticed that spacer RNAs have the same structure and characteristics as miRNAs. Hence, we proposed a simple “reverse activation” strategy (Figure 1c) to directly detect miRNA. According to the design, in the presence of both scaffold RNA and excess ssDNA activators (containing 20-nt deoxyribonucleotides complementary to miRNA), the miRNA target itself instead of spacer RNA unlocks the *trans*-cleavage activity of Cas12a. This design enables direct detection of miRNA without the need for additional reverse transcription and sample amplification steps. Then, we tested to check the sensitivity of our SCas12a fluorescence assay. For this, we designed multiple ssDNA activators targeting the clinically relevant miRNAs that play important roles in cancer progression and are overexpressed in tumour tissues, including miR-21, miR-31, miR-17 and miR-let-7a. For example, we created dilutions of miR-21 targets (Figure 4a) ranging from 10 fM to 1 nM and tested for the detection of each target by addition of excess ssDNA activators, achieving a limit of detection (LoD) of 100 fM. Similarly, we obtained LoDs of 100 fM to 400 fM for the other three miRNAs (Figure S3), indicating that our assay has a good repeatability for detecting different miRNA substrates.

**Figure 4.**
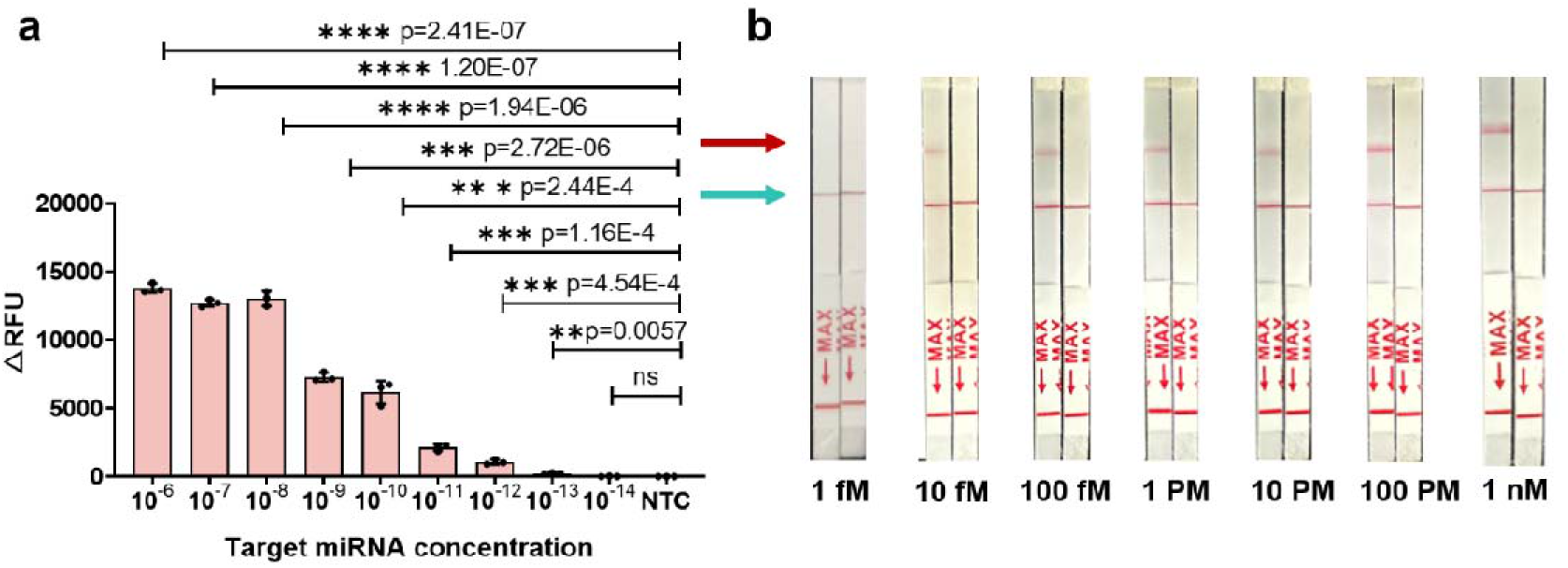
Determination of the limit of detection of the SCas12a-based assays. (a) Limit of detection of the miR-21 target was determined by a fluorescence assay. Plot represents the background subtracted fluorescence intensity at t =10 min, for different concentrations of the target. All error bars represent the mean value +/− SD (n=3) and the statistical analysis was performed using a two-tailed t-test where ns=not significant with p>0.05, and the asterisks (*p≤0.05, **p≤0.01, ***p≤0.001, and ****p≤0.0001) denote significant differences. (b) Similar experiments to (a), except that a dipstick reporter was used instead after the 20 minutes of cleavage reaction. The red arrow indicates the test bands, while the green arrow indicates the control bands.

Next, we measured the sensitivity of the SCas12a system by using a rapid lateral flow assay (Figure 4b). Surprisingly, ten times lower LoD of ∼10 fM was observed. This might be explained by the local concentrations of the components, including Cas12a enzyme, scaffold RNA, ssDNA activator, ssDNA probes and the miRNA targets, were significantly increased in the dipstick. To further enhance the sensitivity of SCas12a system, we are developing an amplification-free digital miRNA assay in combination with a micro fluidic device.

### Application of SCas12a-based fluorescence assay for multifunctional RNA detection

To evaluate the selectivity of our assay, we tested and compared several types of miRNAs (miR-17, miR-31, and miR-let-7a) with the miR-21 target. Compared with miR-21 (Figure 5a), none of the non-targeted miRNAs with 100-fold concentrations enhanced the fluorescence signal, indicating high specificity of SCas12a assay. Next, we tested the ability of our method for multiplexed detection of miRNAs. To investigate this, we used two different ssDNA activators with unique targets (miR-17 and miR-31), or pooled them together. The results (Figure 5b) indicated that the targeted miRNAs can be individually or simultaneously detected without interference by each other. Moreover, SCas12 assay is feasible for multiplexed detection of DNA and RNA when introducing DNA targets and the corresponding spacer RNA activators. This observation is critical because muti-target biomarkers, such as miRNA and circulating tumor DNA (ctDNA), are more accurate for the diagnosis of cancer early than a single indicator. To sum up, our assay has the potential utility for multiplexed miRNA and ctDNA liquid biopsy in cancer diagnosis and prognosis.

**Figure 5.**
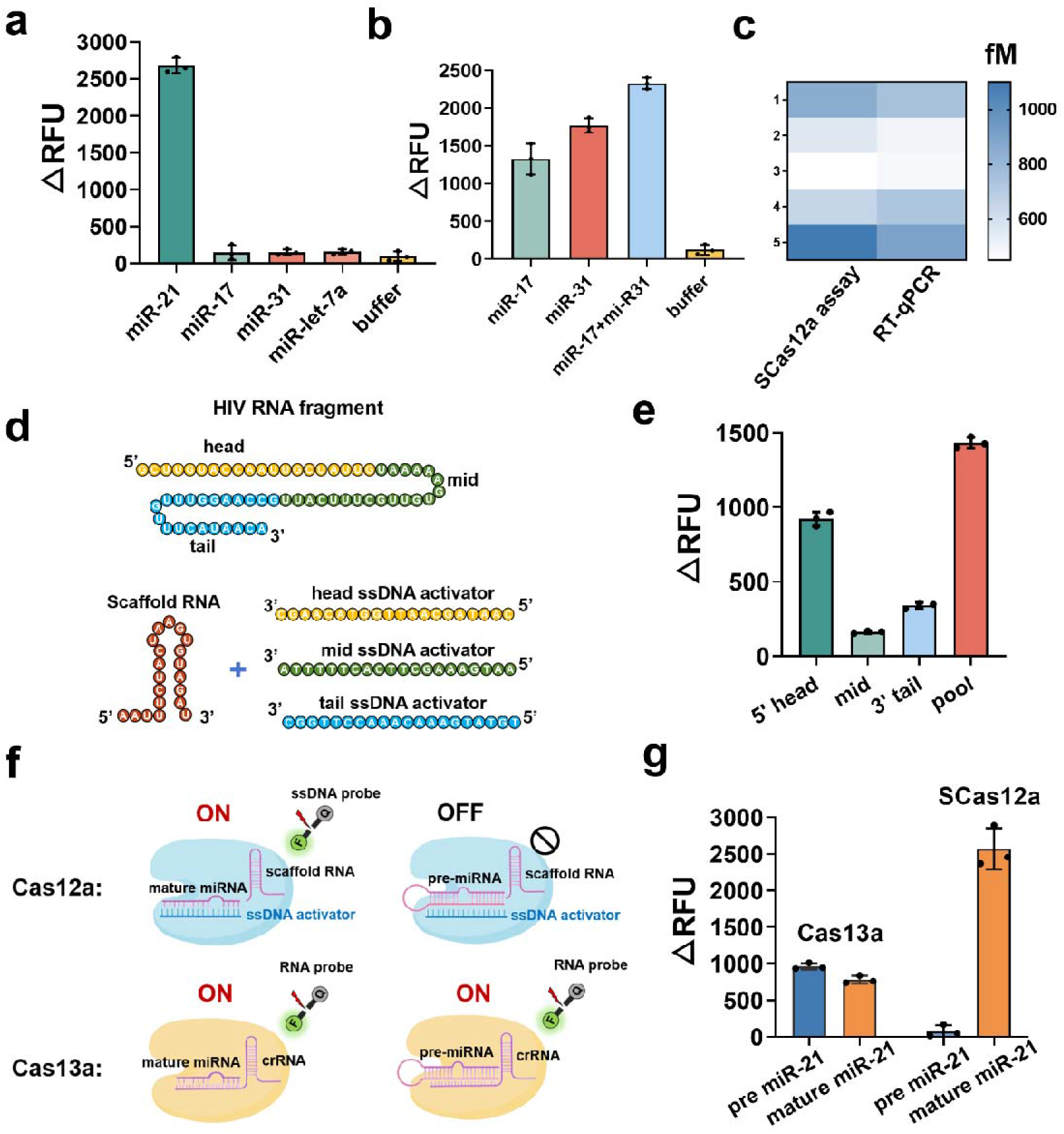
Quantitative detection of RNA by SCas12a assay. (a) Selective detection of the target miR-21 (10 pM) in the different types of miRNAs (1 nM), including miR-17, miR-31 and miR-let-7a. (b) Multiplexed detection of miR-17 and miR-31 in the same reaction. (c) Heatmap of the estimated target miR-21 concentrations of the SCas12a assay and conventional RT-qPCR assay. Schematic of an HIV RNA target and three ssDNA activators targeting it at different positions. Comparison among the 5’ head, mid, 3’ tail and pooled HIV targeting ssDNA activators. (f) Schematic of the detection of mature miRNA and pre-miRNA by SCas12a or Cas13a based fluorescence assay. (g) Comparison of fluorescence intensity changes between pre miR-21 and mature miR-21 detected by Cas13a or SCas12a.

Next, we explored the utility of developed SCas12a assay for measuring miRNAs in real biological samples. We firstly extracted the total RNA from different batches of breast cancer cells (MCF-7) using a commercial RNA extraction kit. Then, we measured miR-17 levels in real-time by SCas12a assay or RT-qPCR. The results (Figure 5c) showed that the two methods have an excellent correlation. In conclusion, we provided a simple, highly sensitive and specific SCas12a-based amplification-free diagnostic method for miRNA detection.

Upon closer inspection of the structure of SCas12a RNP (Figure 1b), we hypothesized that the RNA targets with a high amount of secondary structure, especially in the 5’ end, are more inaccessible to bind to the “Cas12a-scaffold RNA” complex, and are therefore harder to detect, while targets with relatively low or no secondary structure are detected easily. To test this, we first detected a 60-nt RNA fragment of human immunodeficiency virus (HIV) with three designed ssDNA activators (Figure 5e), resulting in significant variability in *trans*-cleavage activity. The head-targeting activator showed as much as a 4 to 6-fold increase in activity as compared to mid-targeting and the tail-targeting activator. Interestingly, the method displayed a higher activity with pooled ssDNA activators. This enhancement of the detection level by pooling together multiple guides has previously been shown with the Cas13a enzyme (18). Therefore, we suggest that the pooled ssDNA activators should be adopted to detect the long-stranded RNA targets.

Finally, we hypothesized the steric-hinerance effect observed above may be utilized to discriminate between mature miRNAs and pre-miRNAs comprising the same miRNA sequence. To verify this, we measured miR-21 and pre miR-21 at the same concentration by SCas12a or Cas13a based fluorescence assays. Using SCas12a (Figure 5f), the mature miR-21 showed 2500-fold fluorescence intensity of pre miR-21. However, using Cas13a (Figure 5f), we did not observe a significant difference in fluorescence intensity between mature miR-21 and pre miR-21. These results suggested false positive results with high levels of mature miRNAs identified by Cas13a-based diagnostic methods, a phenomenon that was most likely overlooked in previous studies. Given that the SCas12a uses a cheaper DNA probe and has 1000 times the sensitivity (10 fM *vs* 10 pM) compared to Cas13a, we believe that it is a better diagnostic method to detect mature miRNAs with high accuracy, sensitivity, and low cost.

### SCas12a-based assay improves mutation specificity of DNA detection

To be a fully assembled RNP (Figure 3), we have shown that the Cas12a protein needs to properly bind to both scaffold RNA and spacer RNA. Given this, we hypothesized that SCas12a might be more sensitive in discriminating DNA single point mutations than wild-type Cas12a. To test this, we designed a 20 nt spacer RNA with single-point mutations for SCas12a detection, as well as a 40-nt crRNA with identical mutations for WT Cas12a detection. The results of a side-by-side comparison were shown in Figure 6. In short, we found that the detection of single-point mutants with SCas12a is more sensitive. In particular, we observed that mutations in position M2-M6, M14 and M19 significantly decreased the SCas12a-mediated *trans*-cleavage activity, while only M4 and M19 mutations obviously decreased the WT Cas12a-mediated *trans*-cleavage activity. Our data indicated that SCas12a has an improved specificity for mutations closer to the 5’ or 3’ ends of spacer RNA. This might be explained by the boundary sequence of the spacer RNA has a great effect on the assembly of SCas12a RNP. Thus, mutations in these positions significantly reduced RNP’s *trans*-cleavage activity. This feature of SCas12a could be utilized to develop robust SNP diagnostics in the future.

**Figure 6.**
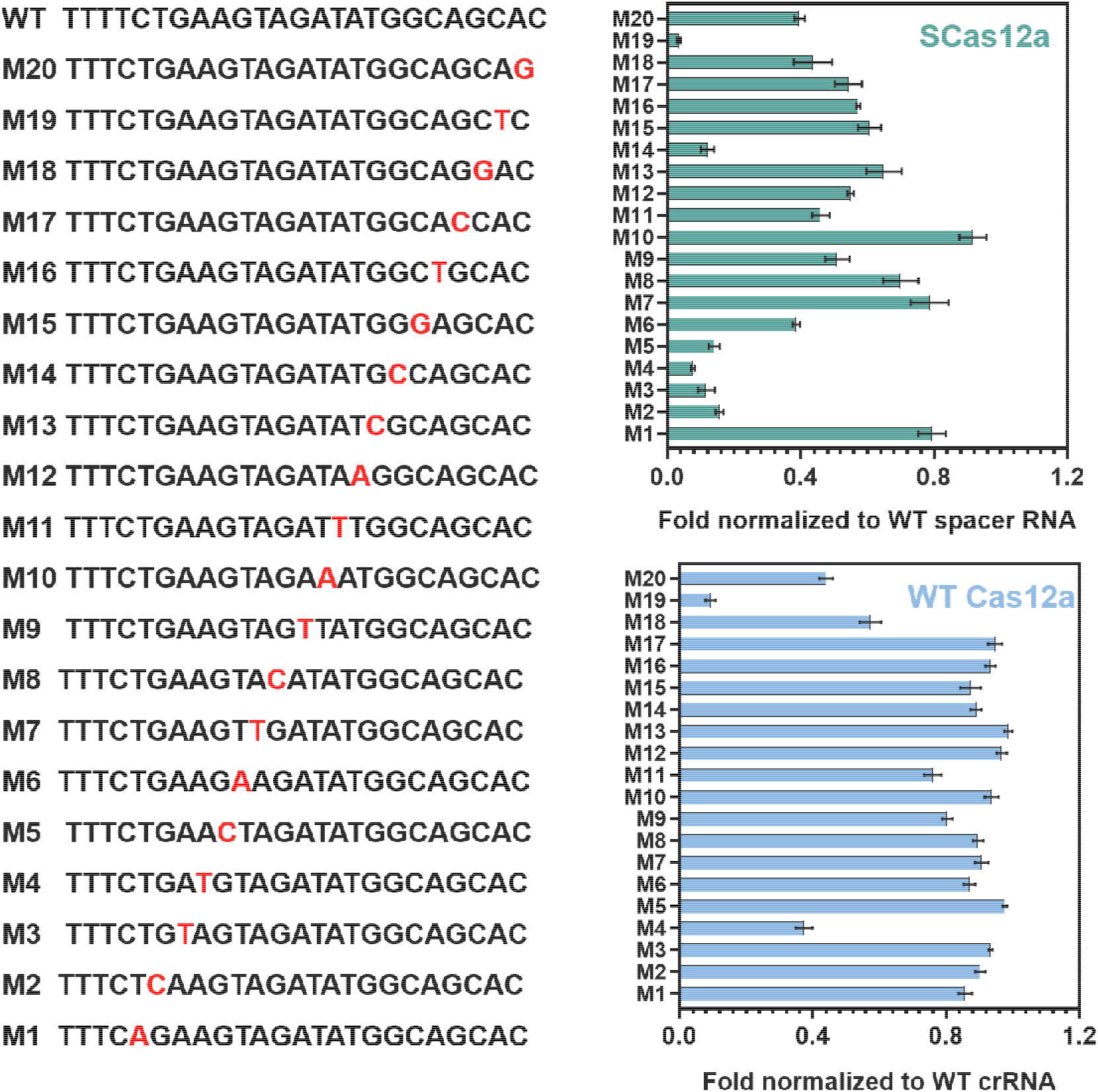
Specificity of SCas12a assay towards single point mutations in DNA target. Spacer RNA (SCas12a) or crRNA (wt Cas12a) were designed with point mutations across the length of the pairing region of a DNA target which derived from HPV16. The mutation location is identified by ‘M’ following the nucleotide number where the base has been changed to its complementary deoxynucleotide (3’ to 5’ direction).

### Application of SCas12a-based assay for DNA detection

WT CRISPR-Cas12a system has been extensively adopted to detect DNA target in combination with nucleic acid amplification means such as polymerase chain reaction (PCR), recombinase polymerase amplification (RPA) (23, 24), loop-mediated isothermal amplification (LAMP) (25, 26), etc. Here, we firstly tested the sensitivity of SCas12a assay for amplification-free detection of DNA. As a case study, we chose human papillomavirus (HPV) 16 as the test DNA substrate. Briefly, HPV16-containing plasmids were incubated with ssDNA fluorescence probes and the SCas12a RNP targeting HPV16 fragments. We obtained a LoD of 10 pM by fluorescence assay (Figure 7a), which was consistent with the previous results obtained by WT Cas12a (17). In addition, a LoD of 1 pM was achieved by rapid lateral flow assay (Figure 7b). To further improve the sensitivity of the method, we developed a one-pot method combining RPA isothermal amplification with fluorescence analysis to achieve the target identification at attomolar levels (Figure 7a).

**Figure 7.**
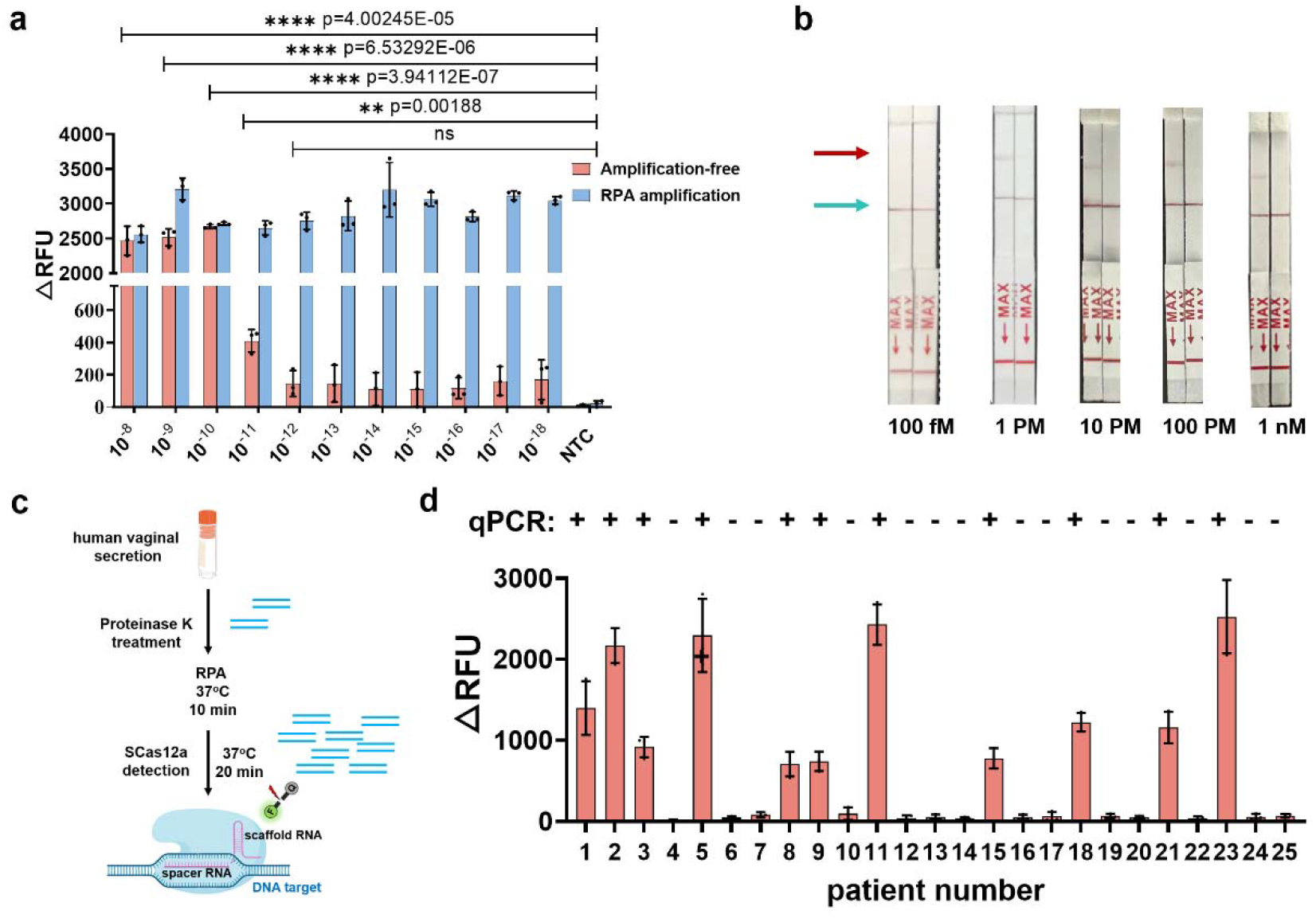
Rapid detection of HPV16 by SCas12a assay. (a) Determination of limit of detection of DNA by SCas12a fluorescence assay with or without recombinase polymerase amplification (RPA). (b) Direct detection of DNA by SCas12a-based lateral flow assay. (c) Schematic outlining DNA extraction from human vaginal secretion samples to HPV identification by SCas12a fluorescence assay. (d) Identification of HPV16 in 25 patient samples by PCR (up) and SCas12a fluorescence assay (down).

To investigate the utility of our method for analyzing patient samples, we tested crude DNA extraction from 25 human virginal secretion samples (Figure 7c) that had previously been analyzed for HPV16 infection by qPCR. Within 30 minutes, the method accurately identified HPV16 (25/25 agreement) in patient samples, showing excellent correlation with qPCR results (Figure 7d). Our data also demonstrated that the one-pot SCas12a assay has the same DNA detection capabilities as DETECTER (3). Collectively, we offer a new SCas12a-based diagnostic method for rapid, cost-effective, and highly sensitive DNA testing.

## Discussion

CRISPR-Cas12a has been extensively studied and widely used in DNA-based diagnostic platforms. When detecting RNA by Cas12a, reverse transcription and subsequent amplification reactions are required since the Cas12a enzyme does not naturally tolerate RNA substrates. Recently, Jain et al. developed a split-activator-based method called SAHARA (19) for programmable RNA detection with Cas12a. However, their diagnostic method has two major limitations: (i) only picomolar levels (250-700 pM) of RNA detection was obtained which does not meet the needs of clinical testing, and (ii) the specificity in more complex samples is not good enough since the method is designed to only bind to 12-nt of the target RNA. Upon deep inspection of the “Cas12a-crRNA-split activator” complex they obtained, we found that this complex is too crowded to make it difficult for the RNA substrates to enter, and therefore resulted in decreased *trans*-cleavage activity. To resolve these issues, here we proposed a simpler strategy by combination Cas12a enzyme with a split crRNA that was divided into scaffold RNA and spacer RNA. Compared to the wild-type Cas12a, the structure changes of the designed SCas12a RNP are minimal.

In this work, we first found that the SCas12a RNP has a *trans*-cleavage activity comparable to that of wild-type Cas12a RNP. Taken advantage of miRNA targets as the spacer RNA and another ssDNA as the activator, we devised SCas12a-based fluorescence and rapid lateral flow assays, achieving highly sensitive (LoD of 10 fM) and selectively detection of miRNA without the need for reverse transcription and pre-amplification. In addition, our assay enabled multiplexed detection of different miRNAs, quantitative analysis of miRNAs from cancer cells, as well as identification of long-stranded RNAs without secondary structure. Compared to conventional Cas13a-based diagnostic methods, this SCas12a assay can discriminate between mature miRNA and pre-miRNA that contains the same sequence of miRNA. Moreover, we showed that SCas12a outperforms wild-type Cas12a in specificity towards DNA point mutations, a feature that can be used to develop robust SNP diagnostics. Finally, we combined RPA with fluorescence method to establish a one-pot SCas12a detection platform, and realized the detection of attomolar concentrations of HPV16 in patient samples.

In summary, we developed a SCas12a-based diagnostic method with major advantages: (1) much cheaper and more stable 20-nt spacer RNA instead of 40-nt crRNA was adopted (2) mature miRNA rather than pre-miRNA was accurately identified for the first time; (3) multiple cancer biomarkers such as miRNA and ctDNA can be simultaneously detected (4) the specificity towards point mutations is higher than WT Cas12a-based methods. Furthermore, the simple “split-crRNA-activator” strategy of this work can be expanded to other CRISPR-Cas enzymes for developing innovative molecular diagnostics. In conclusion, this work presents a powerful nucleic acid detection approach with the potential for commercial use in clinical applications.

## Materials and Methods

### Ethical statement

For this study, human virginal section samples for HPV16 detection were collected and provided by the department of pharmacy in Henan cancer hospital with a protocol approved by the ethics committee at Zhengzhou University.

### Materials

All DNA, RNA, FAM-labeled ssDNA, and FAM-labeled RNA fluorescence reporter that were listed in supplementary Table S1 were ordered from Sangong Biotech (Shanghai, China). The general chemicals for buffer preparation were from Sinopharm (Beijing, China). The 10xNEB buffer2.1 and 10xCutsmart buffer were purchased from New England Biolabs (Beijing, China). The proteinase K, and RNA extraction kit were bought from Takara (Dalian, China). Ultrapure water was employed throughout the work. All DNA oligonucleotides and RNA were dissolved in the DEPC (RNase-free) water, and stored at −20°C for subsequent experiments. A fluorescence spectrophotometer F4600 (Hitachi, Japan) and a CFX96 real-time system (Bio-Rad, USA) were used for the fluorescence measurements. The excitation and emission wavelengths of FAM were 485 nm and 520 nm, respectively.

### Recombinant AsCas12a and LwbCas13a protein purification

The synthesized AsCas12a (*Acidaminococcus sp*. Cas12a) gene or LwaCas13a (*Leptotrichia wadei*. Cas13a) was inserted into pET-28a(+) vector. Next, the correctly constructed expression plasmids were transformed into *Escherichia coli* BL21 (DE3) competent cells. Following a standard purification protocol (27), the recombinant AsCas12a proteins were purified by running a Ni-NTA column and a HiLoad® 16/600 Superdex® size-exclusion column. Finally, the purified AsCas12a and LwbCas13 proteins were concentrated using Millipore concentrators (100 KD), analyzed by 10% SDS-PAGE (Figure S1), and snap-frozen in liquid nitrogen, and stored at –80°C until use.

### Polyacrylamide gel electrophoresis

Firstly, the *cis*-cleavage and *trans*-cleavage activity of WT Cas12a and SCas12a RNP were determined by a standard protocol (27). Next, the reaction products were resolved on 12% polyacrylamide gel using 1X TBE as a running buffer at a constant voltage of 100 V for 120 min. After staining with GoldView, the gel image was taken with an UV transilluminator.

### Conventional Cas12a-based or Cas13a-based fluorescence assay

The cleavage reaction was conducted in a final volume of 20 µL, including 2 μL 10X NEBuffer 2.1 buffer, 0.5 μL AsCas12a (1 μM) or LbCas13a (1 μM), 1 μL crRNA (1 μM), 1 μL fluorescence-quencher probe (10 μM), 13.5 μL RNase-free water and 2 μL various concentrations of target ssDNA (Cas12a) or miRNA (Cas13a). The fluorescence signal was measured at 30 s intervals at 37°C using a CFX96 touch real-time PCR system (Bio-Rad, CA, USA).

### Application of SCas12a assay for RNA detection

All fluorescence-based detection assays were carried out in a CFX96 touch real-time PCR system (Bio-Rad, CA, USA). The Cas12a, scaffold RNA and ssDNA activators were assembled by mixing them in NEB buffer 2.1 and nuclease-free water followed by incubation at room temperature for 10 min. Then, the mixtures were added to 1000 nM FQ reporter and the necessary concentration of the target miRNA in a 20 μL reaction volume. The reactions were conducted at 37°C for 20 minutes and the fluorescence changes at 520 nm were recorded. A final concentration of 250 nM Cas12a, 500 nM scaffold RNA, and 500 nM of target ssDNA activators were used in all the assays unless otherwise specified. The lateral flow assays were performed in the same reactions except using a dipstick containing ssDNA probes for cutting.

### RNA extraction and RT-qPCR

Total RNA was extracted from MCF-7 cancer cells by using the RNA extraction kit from GenePharma (Suzhou, China). Next, the cDNA was synthesized from the extracted total RNA using the miRCURY LNA RT Kit (Qiagen). Briefly, a reverse transcription reaction was performed at 42°C for 60 min and then inactivated at 95°C for 5 min. Synthesized cDNA was stored at −20°C before use. For detection of specific miRNA targets, the stem-loop primers and Hairpin-it miRNA RT-qPCR detection kit were designed and ordered from GenePharma (Suzhou, China). Then, we quantitatively measured the miRNA targets in cells following our reported protocol (20).

### Application of SCas12a assay for DNA detection

Direct detection of DNA targets by SCas12a-based fluorescence assay or lateral flow assay was carried out following the same protocol as mentioned above for RNA detection except for spacer RNA activators were used instead of ssDNA activators. To enhance the sensitivity of detection, a recombinase Polymerase Amplification (RPA) using TwistAmp Basic (TwistDx) was performed followed by SCas12a detection in the same reaction. Briefly, 40 µl reactions containing 1 µl sample, 0.4 µM forward and reverse primer, 1× rehydration buffer, 14 mM magnesium acetate and RPA mix were incubated at 37°C for 10 minutes. Then, the RPA reaction was transferred to the PCR tube containing Cas12a, scaffold RNA, spacer RNA, and FQ reporter in a 20 μL reaction volume. Finally, the reactions were conducted at 37°C for 20 minutes and the fluorescence changes at 520 nm were recorded. For HPV identification by SCas12a fluorescence assay, detection values of HPV16 human samples were normalized to the maximum mean fluorescence signal obtained using spacer RNA targeting the hypervariable loop V of the L1 gene within HPV16 (28). A one-way ANOVA with Dunnett’s post-test was used to determine the positive cutoff (set at p ≤ 0.05) for identification of HPV16 in patient samples. Based on this cutoff, 100% of samples were accurately identified for HPV16 infection (25/25 agreement with PCR-based results.

## Data Availability

All other study data are included in the article and/or SI Appendix.

## Acknowledgments

This work was supported by the National Key Research and Development Program of China (2022YFC2304304), National Natural Science Foundation of China (22007030), Science and Technology Innovation Talent Plan od Hubei Province (2023DJC136), the Open Funding Project of the State Key Laboratory of Esophageal Cancer Prevention (K2022-008), the Open Funding Project of the State Key Laboratory of Biocatalysis and Enzyme Engineering, and Research and Innovation Initiatives of WHPU (2022J02).

---

**Figure S1.**
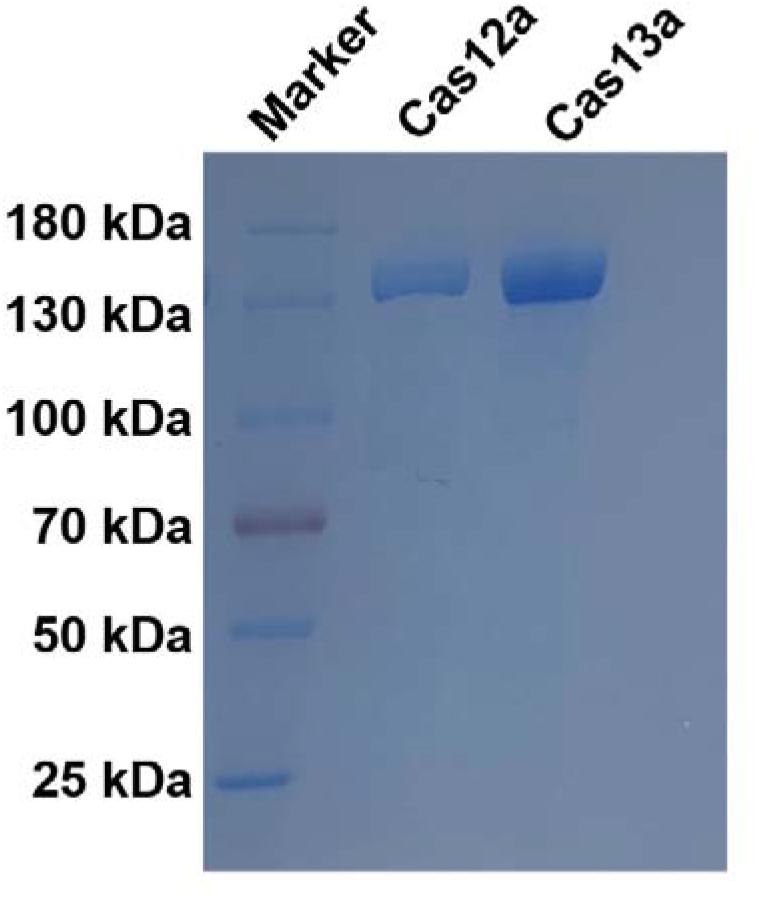
10 % SDS-PAGE analysis of the purified recombinant AsCas12a and LwaCas13a proteins.

**Figure S2.**
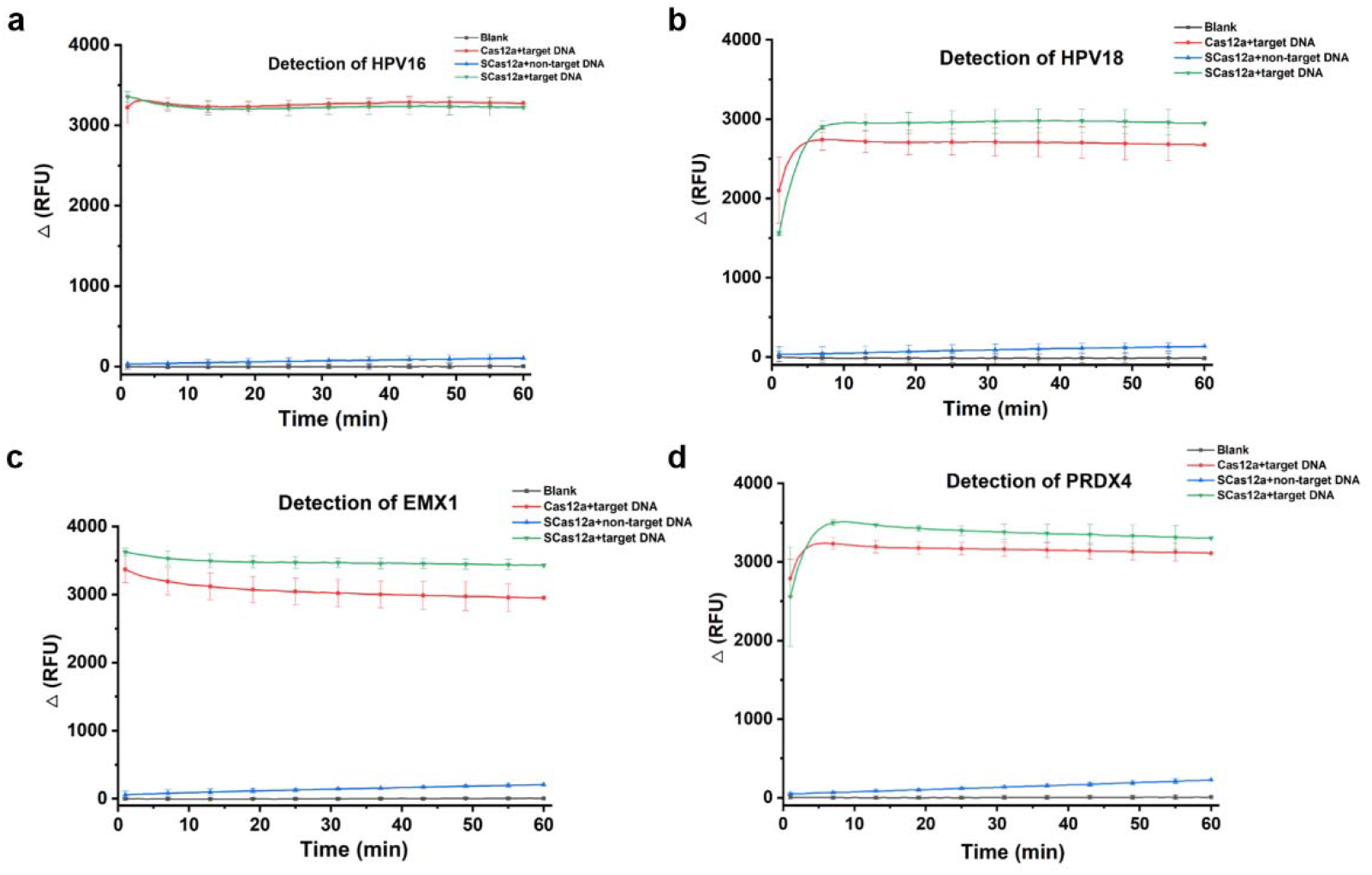
The real time fluorescence recording for the *trans*-cleavage of different targeted DNAs by wild-type Cas12a or SCas12a, including the gene fragments derived from (a) HPV16, (b) HPV18, (c) EMX1 and (d) PRDX4.

**Figure S3.**
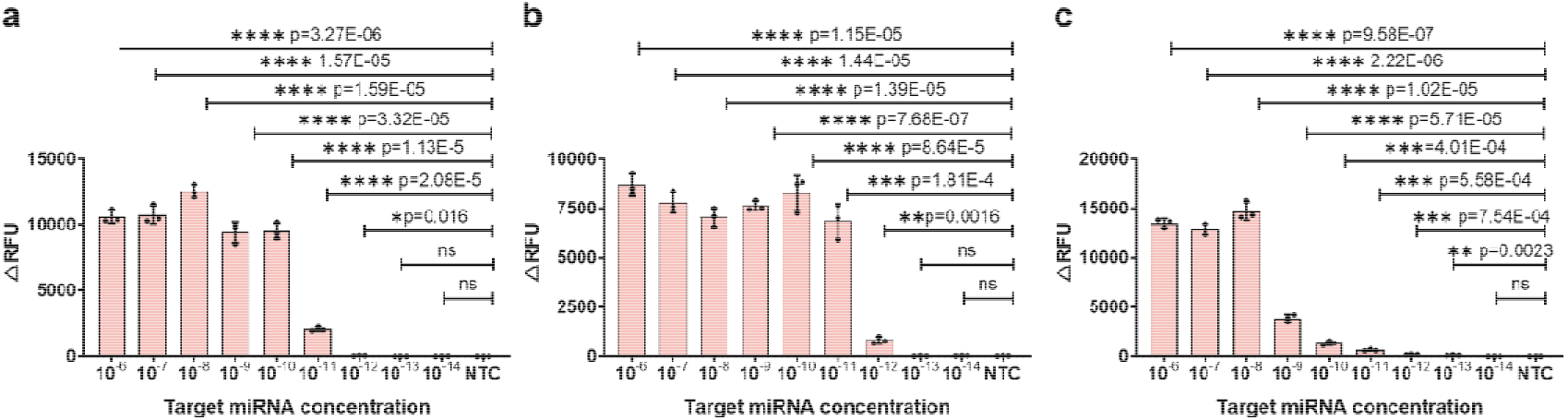
Determination of the limit of detection of SCas12a fluorescence assay using different miRNA substrates, including (a) miR-17, (b) miR-31, and (c) miR-let-7a.

**Table S1.**
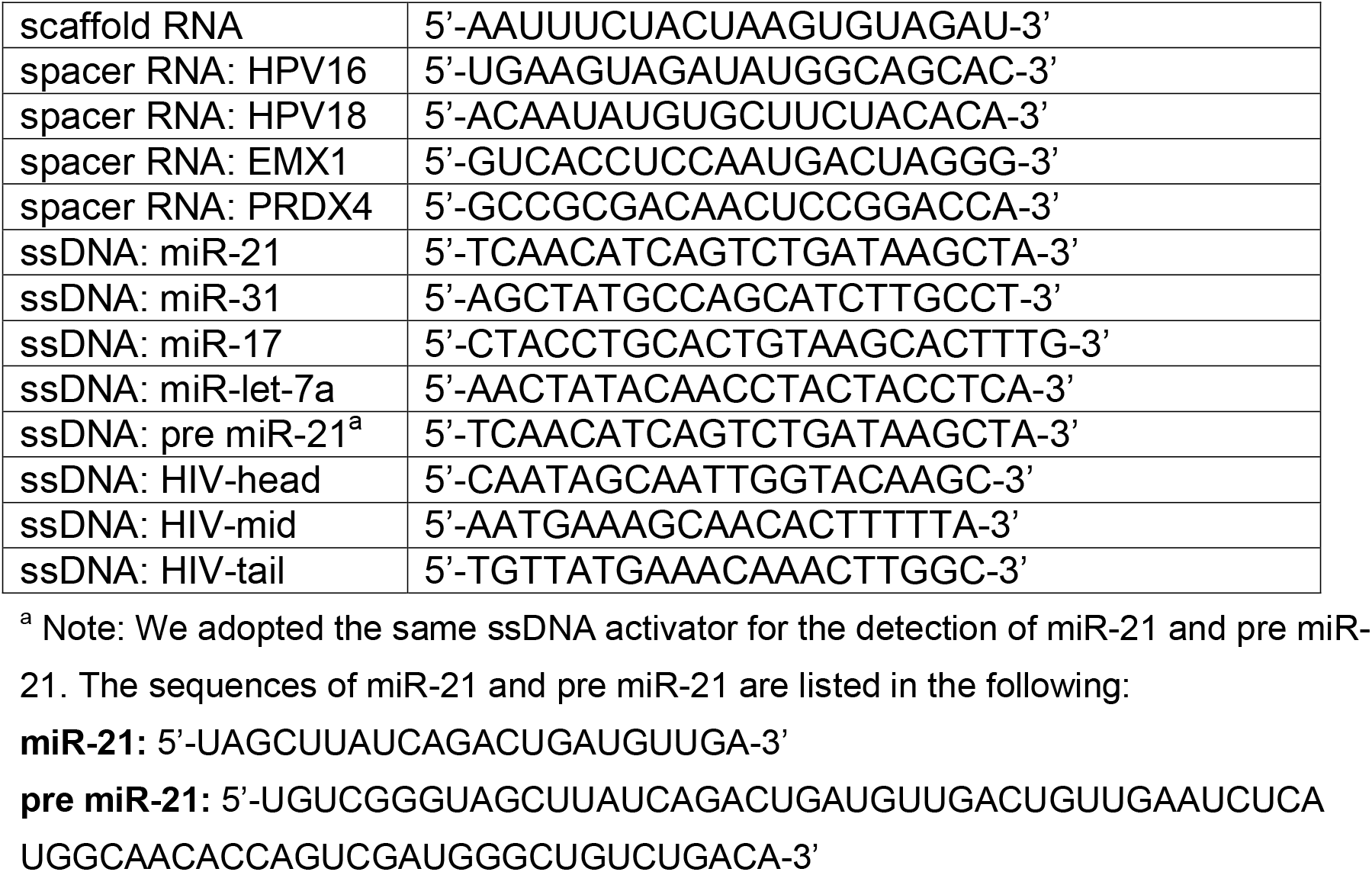
Sequences of the RNAs and ssDNA activators used in this work.

